# Predictive Neuromechanical Simulation Explains Gait Biomechanics in Obesity

**DOI:** 10.64898/2026.06.03.729794

**Authors:** Chi-Whan Choi, Vincent Ton, Simone V. Gill, Seungmoon Song

**Affiliations:** Sargent College of Health and Rehabilitation Sciences, Boston University, Boston, MA, USA; Department of Mechanical and Industrial Engineering, Northeastern University, Boston, MA, USA

**Author notes:** **Corresponding Author:** Dr. Chi-Whan Choi, 111 Cummington Mall, Boston MA, 02215. E-mail addresses (V. Ton), (S. V. Gill), and (S. Song).

**Keywords:** **Keyword:** Obesity, Neuromechanical simulations, Gait biomechanics, Knee joint loading

## Abstract

Individuals with obesity exhibit gait adaptations including reduced early-stance knee flexion, altered muscle coordination, slower preferred walking speeds, and shorter step lengths. Although these features are well documented, the mechanisms by which obesity-related physiological changes produce these patterns and influence knee joint loading relevant to osteoarthritis (OA) remain unclear. This study used predictive neuromechanical simulation to examine how musculoskeletal changes and movement objectives interact to generate obesity-associated gait patterns and tibiofemoral loading. Predictive simulations were performed using a reflex-based neuromechanical walking model. A baseline non-obese model (1.8 m, 80 kg) was modified to represent obesity-related changes in segment mass distribution and muscle strength (1.8 m, 140 kg), including more apple-like and more pear-like body mass distributions. Control parameters were optimized to generate stable walking while minimizing muscle effort and tibiofemoral joint loading. Objective weightings were identified by matching simulated knee kinematics to experimental observations at a typical walking speed. Using the selected weightings, we compared joint kinematics, kinetics, and muscle activations, and simulations were performed across walking speeds to evaluate optimal walking speed, step length, muscle effort, and knee loading. The baseline model best matched reference knee kinematics using a muscle-effort objective alone, whereas the obese model required a combined objective penalizing both muscle effort and knee loading. This formulation reproduced key gait features, including reduced early-stance knee flexion, reduced vastii activation with increased plantarflexor activation, slower optimal walking speeds, and shorter step lengths. Variations in body mass distribution produced moderate but consistent effects on gait mechanics relative to larger effects of increased body mass. Obesity-related changes in body mass and muscle strength alone did not reproduce observed gait patterns, but incorporating an objective that penalizes knee loading generated multiple characteristic features. Predictive neuromechanical simulation provides a framework for identifying candidate mechanisms linking obesity, gait biomechanics, and knee joint loading.

**Author Summary:** Understanding how and why obesity alters gait is a complex biomechanical problem involving multiple interacting factors including increased segmental mass, altered inertial properties, and reduced relative muscle strength. These factors interact in ways that are difficult to isolate through experimental observation alone. Here, we used computer simulations to examine how musculoskeletal changes and movement objectives interact to generate obesity-associated gait patterns and knee loading. We found that physiological changes alone did not reproduce observed gait features, whereas incorporating an objective that penalizes knee loading generated multiple characteristic features simultaneously, including reduced early-stance knee flexion, altered muscle coordination, slower optimal walking speeds, and shorter step lengths. These findings suggest that obesity-associated gait reflects coordination strategies that regulate knee loading under increased body mass.

## Introduction

Understanding how and why obesity alters gait could inform strategies to promote physical activity and reduce the risk of movement-related musculoskeletal conditions in individuals with obesity. An increase in mechanical loading caused by greater body weight is a likely major contributor to musculoskeletal problems such as knee osteoarthritis (OA) [1,2], but individuals with obesity also commonly exhibit altered gait patterns that may further influence how forces are distributed across the lower-limb joints [2]. These gait differences raise important unresolved questions about their underlying causes. For example, do they reflect adaptive strategies that help regulate knee loading, or do they arise from physiological constraints such as increased segmental mass, altered inertial properties, or reduced relative muscle strength that could inadvertently affect knee loading [3,4]? Substantial inter-individual variability further complicates interpretation, because obesity is associated with multiple musculoskeletal and gait characteristics reported in prior literature [5–14] (Table 1), including differences in body mass distribution such as more apple-like versus more pear-like patterns that are known to affect gait mechanics and muscle activation [5]. Clarifying these mechanisms is essential for determining how obesity influences knee loading and for guiding interventions designed to reduce the risk or progression of knee OA.

**Table 1.**
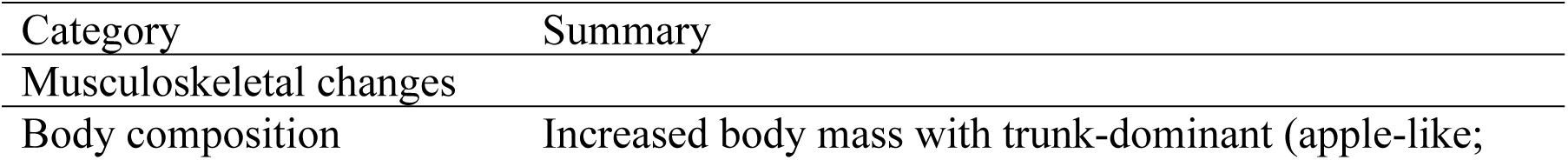
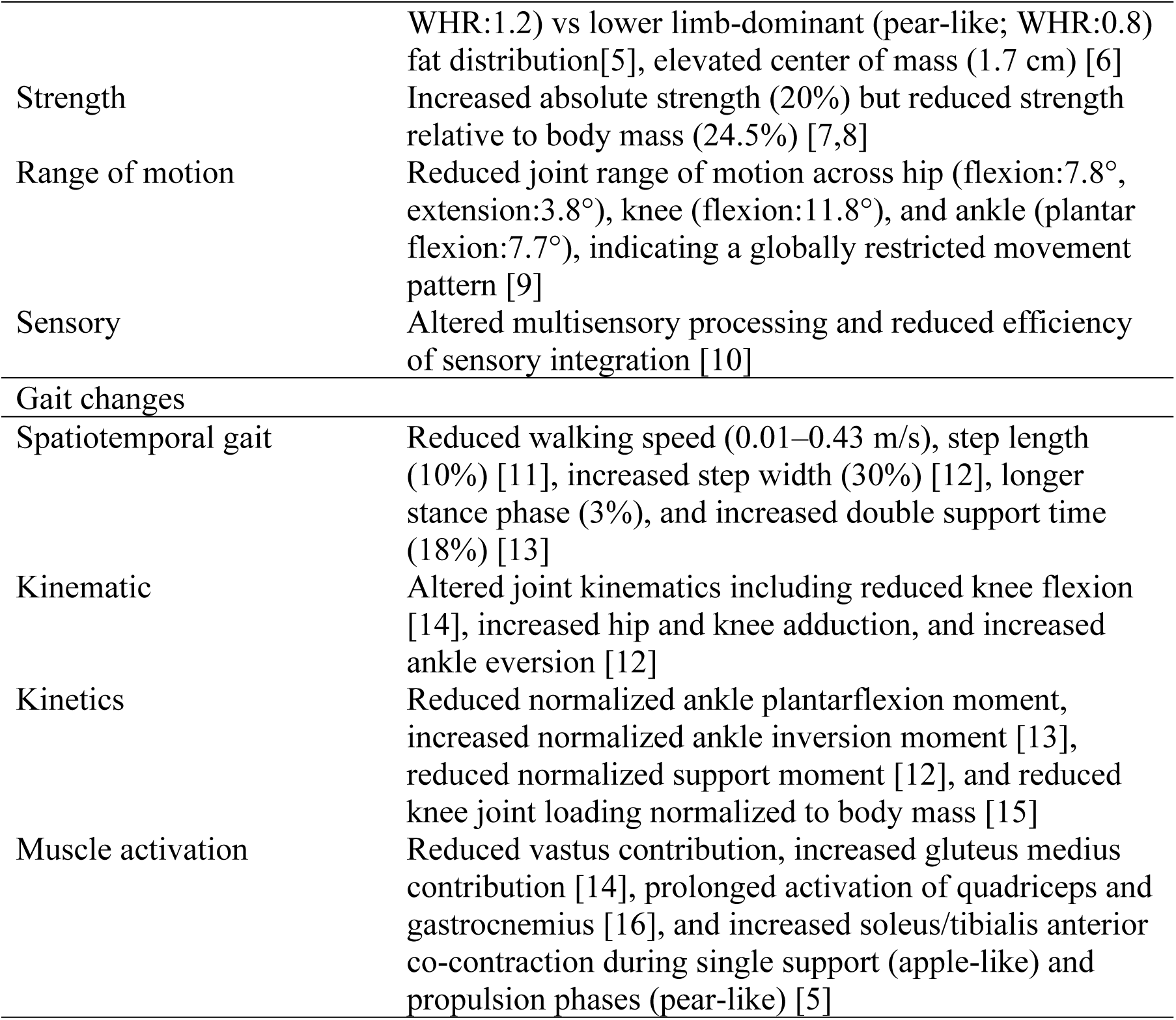
Experimentally reported musculoskeletal and gait characteristics associated with obesity. Summary of physiological and biomechanical features reported in prior literature that inform interpretation of obesity-related gait adaptations. Not all listed characteristics were explicitly represented in the present simulations. WHR: waist-to-hip ratio.

While motion analysis, inverse dynamics, and musculoskeletal tracking simulations have been successfully used to characterize gait in individuals with obesity [14–16], these approaches cannot address key questions about why specific gait patterns emerge and the potential effect of altered gait. Prior studies have described obesity related differences in joint kinematics, joint moments, and muscle activation during walking, including reduced knee flexion in early stance [14], prolonged gastrocnemius and quadriceps activity [16], slower preferred speeds [12], and shorter step lengths [11]. However, because these studies rely on inverse or tracking based analyses of experimentally collected gait data, they cannot determine the underlying reasons for these gait patterns. For example, they cannot explain why individuals with obesity prefer to walk more slowly with shorter steps even though they are capable of walking faster, nor can they determine whether the observed gait characteristics mitigate or worsen knee loading. These limitations highlight the need for modeling approaches that can systematically manipulate physiological properties and gait objectives in order to identify the mechanisms that drive obesity related gait adaptations.

Predictive neuromechanical simulations have the potential to uncover the mechanisms that give rise to specific gait patterns. Rather than reconstructing movement from experimental data, these approaches use a musculoskeletal model actuated by a neuromuscular controller in forward dynamic simulations, and locomotion emerges from optimizing the controller parameters with respect to a prescribed objective function [17–20]. For walking, objective functions are typically constructed based on consensus in human gait biomechanics and often include terms that minimize muscle activation or metabolic energy expenditure, sometimes combined with performance related costs [21]. Predictive simulation has been applied in a variety of contexts, including understanding the biomechanical effects of muscle weakness, altered morphology, impaired neuromuscular control, and gait assistive devices [22–24]. Reflex based controllers, which rely on physiologically interpretable feedback pathways, have been widely adopted because they can generate stable, human like gait across diverse conditions and can shows human like responses to external perturbations [25,26]. We previously used this framework to explain age related gait adaptations by showing that gait patterns in older adults emerged when the model incorporated musculoskeletal physiological changes associated with aging and identified the loss of leg muscle strength, particularly in fast twitch fibers, as a primary contributor to slower preferred walking speeds [27]. This study illustrated how predictive neuromechanical simulations can reveal the interaction between physiological modifications and gait objectives, motivating the use of similar methods to investigate gait adaptations in individuals with obesity.

Here, we used predictive neuromechanical simulation to investigate the mechanisms underlying gait adaptations associated with obesity. We hypothesized that characteristic features of obese gait emerge from the interaction between obesity-related physiological changes and control objectives that regulate both muscle effort and knee loading. Using a reflex-based simulation framework, we examined how these factors influence knee kinematics, muscle demand, and preferred walking speed. We further evaluated how differences in body mass distribution (more apple-like vs more pear-like) affect predicted gait biomechanics. This approach provides a mechanistic framework for understanding how obesity-related changes in physiology and motor control may influence knee joint loading relevant to OA risk.

## Results

### Balancing muscle effort and knee loading reproduces characteristic obese knee kinematics

The best match to reference early-stance knee flexion at 1.5 m/s was obtained using different objective weightings for the baseline and obese models (Fig 1). The baseline model best matched the reference data using the muscle effort cost alone (*w*_ACT_ = 1.0, *w*_KL_ = 0.0) (Fig 1A, 1B), whereas the nominal obese model required inclusion of both muscle effort and knee-loading costs (*w*_ACT_ = 0.6, *w*_KL_ = 0.4) (Fig 1C, 1D). With their respective optimal weight sets, the models captured the experimentally observed reduction in early-stance knee flexion with obesity [15], with maximum knee flexion angles of approximately 25° in the baseline model and 20° in the obese model. (Fig. 1C, black dashed line vs. 1A, black solid line).

**Fig 1.**
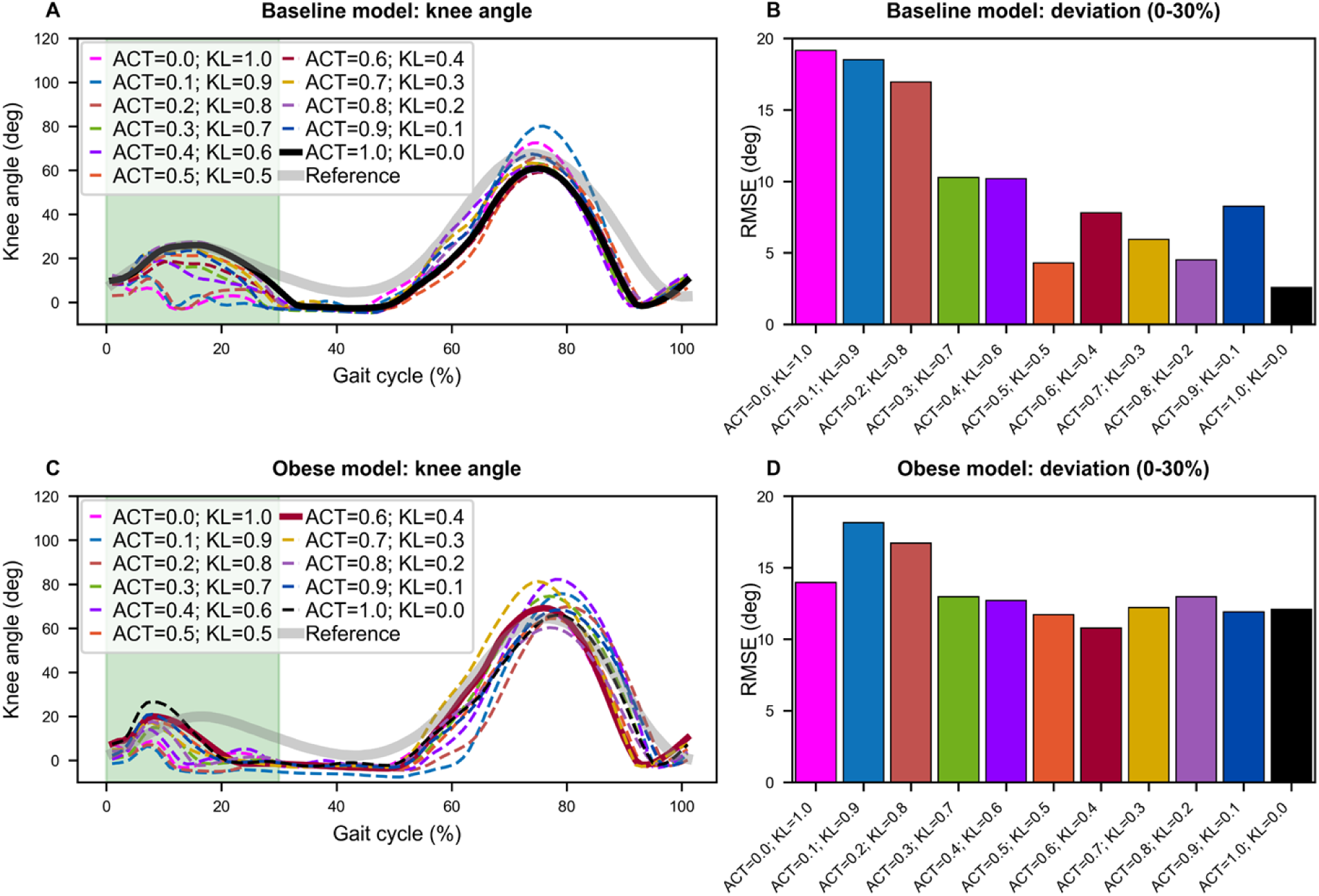
Knee kinematics of the baseline and nominal obese models across cost-term weight combinations at 1.5 m/s. **A**. Baseline model (80 kg): the black solid line shows the knee angle from the weight combination (*w*_ACT_ = 1.0, *w*_KL_ = 0.0) that best matched the non-obese reference data (gray line[15]) during early stance (0–30% of the gait cycle). **B**. RMSE of knee flexion angle during early stance for the baseline model. **C**. Nominal obese model (140 kg): the dark red solid line shows the knee angle from the weight combination (*w*_ACT_ = 0.6, *w*_KL_ = 0.4) that best matched the obese reference data (gray line[15]) during early stance. **D**. RMSE of knee flexion angle during early stance for the obese model. RMSE: root mean squared error.

Compared with the baseline model, the obese model optimized with the combined objective (Obese, ACT+KL, *w*_ACT_ = 0.6, *w*_KL_ = 0.4) best matched early-stance knee flexion exhibited additional gait changes consistent with experimental observations (Fig 2; Supplementary Video). Incorporating the knee-loading cost reduced activation of the vastus muscles associated with early-stance knee flexion, aligning with experimentally observed reductions in knee extensor demand in obesity [14,15]. This reduction in vastus activation was accompanied by increased activation of ankle muscles (i.e., GAS, SOL, and TA), consistent with experimentally reported increases in distal muscle contribution during walking in individuals with obesity [5,14], while absolute knee loading remained greater due to increased body mass. When the obese model was optimized for muscle effort only (Obese, ACT; *w*_ACT_ = 1.0, *w*_KL_ = 0.0), vastus activation increased relative to the combined objective and did not reflect experimentally observed patterns.

**Fig 2.**
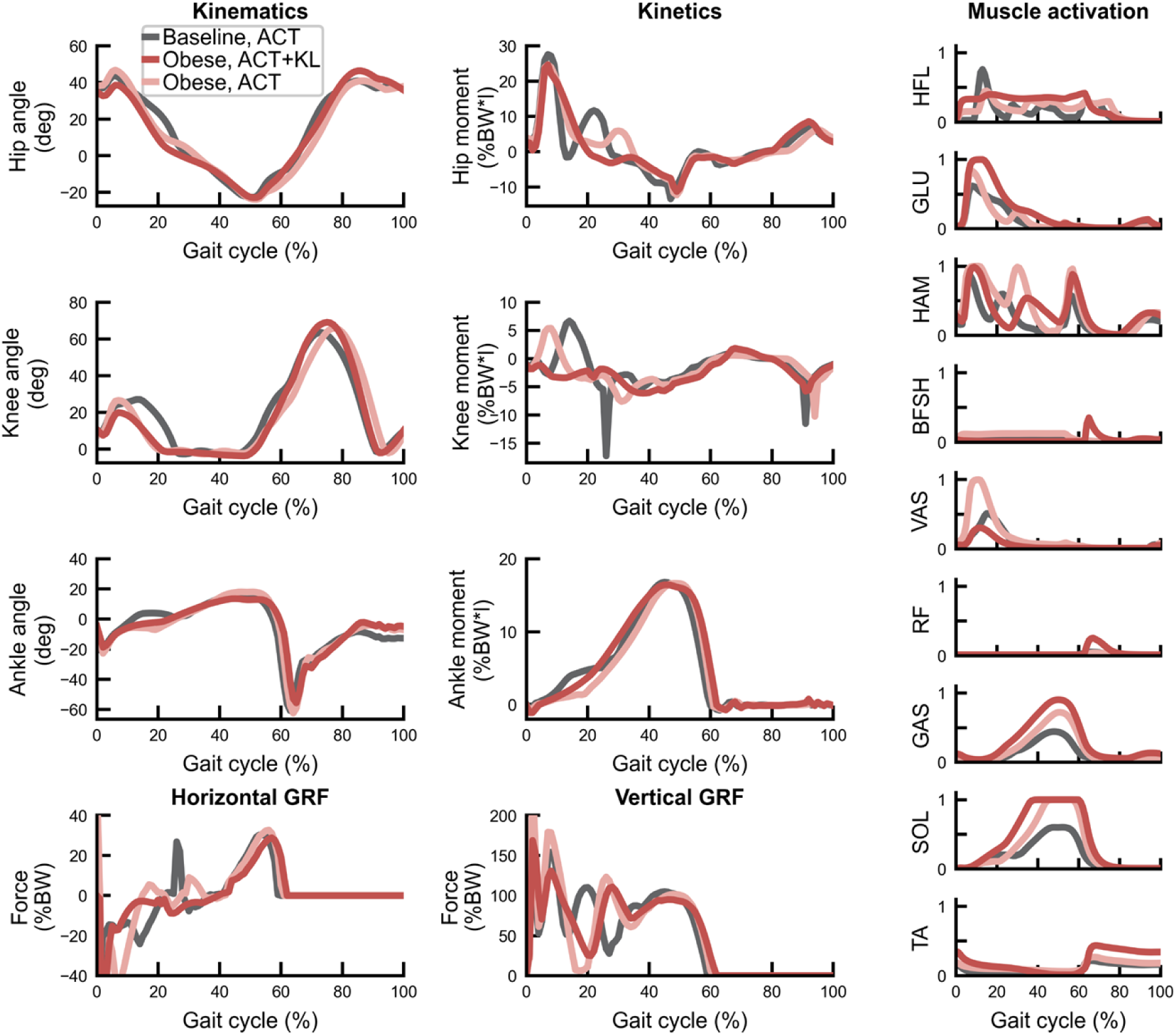
Gait biomechanics at 1.5 m*/*s for Baseline, ACT (*w*_ACT_ = 1.0, *w*_KL_ = 0.0), Obese, ACT (*w*_ACT_ = 1.0, *w*_KL_ = 0.0), and Obese, ACT+KL (*w*_ACT_ = 0.6, *w*_KL_ = 0.4). Joint moments are normalized by body weight (BW) *×* leg length (*l*), ground reaction forces by BW, and muscle activations to a maximum activation of 1.0.

The muscle-effort-only objective (*w*_ACT_ = 1.0, *w*_KL_ = 0.0) is used as the default for the baseline model, and the combined objective (*w*_ACT_ = 0.6, *w*_KL_ = 0.4) for the obese models. Cross-objective simulations are included where helpful for comparison.

### Obese model predicts slower optimal walking speed and shorter step length

Simulations across walking speeds showed that the obese model exhibited slower optimal walking speeds than the baseline model when evaluated using both the muscle effort and knee-loading costs (Fig 3). Across muscle effort, knee loading, and combined cost evaluations (Fig 3A–C), optimal speeds predicted using the selected configurations (*w*_ACT_ = 1.0, *w*_KL_ = 0.0 for Baseline, ACT; *w*_ACT_ = 0.6, *w*_KL_ = 0.4 for Obese, ACT+KL) ranged from 1.26–1.31 m/s for the baseline model and 1.09–1.23 m/s for the obese model, within experimentally observed ranges [12]. The obese model with the selected combined objective consistently predicted lower optimal speeds than the baseline model across all three cost evaluations, with differences of 0.08–0.17 m/s, comparable to reported obesity-related reductions in walking speed [11]. In contrast, when the obese model was optimized using the muscle effort cost alone (*w*_ACT_ = 1.0, *w*_KL_ = 0.0), the predicted optimal speeds were similar to or higher than those of the baseline model depending on the cost term considered (Fig 3D–F). When the baseline model was optimized using the combined objective (*w*_ACT_ = 0.6, *w*_KL_ = 0.4), the optimal speeds deviated from those obtained using the muscle effort cost alone, with the direction and magnitude of change depending on the cost term evaluated (Fig 3G–I).

**Fig 3.**
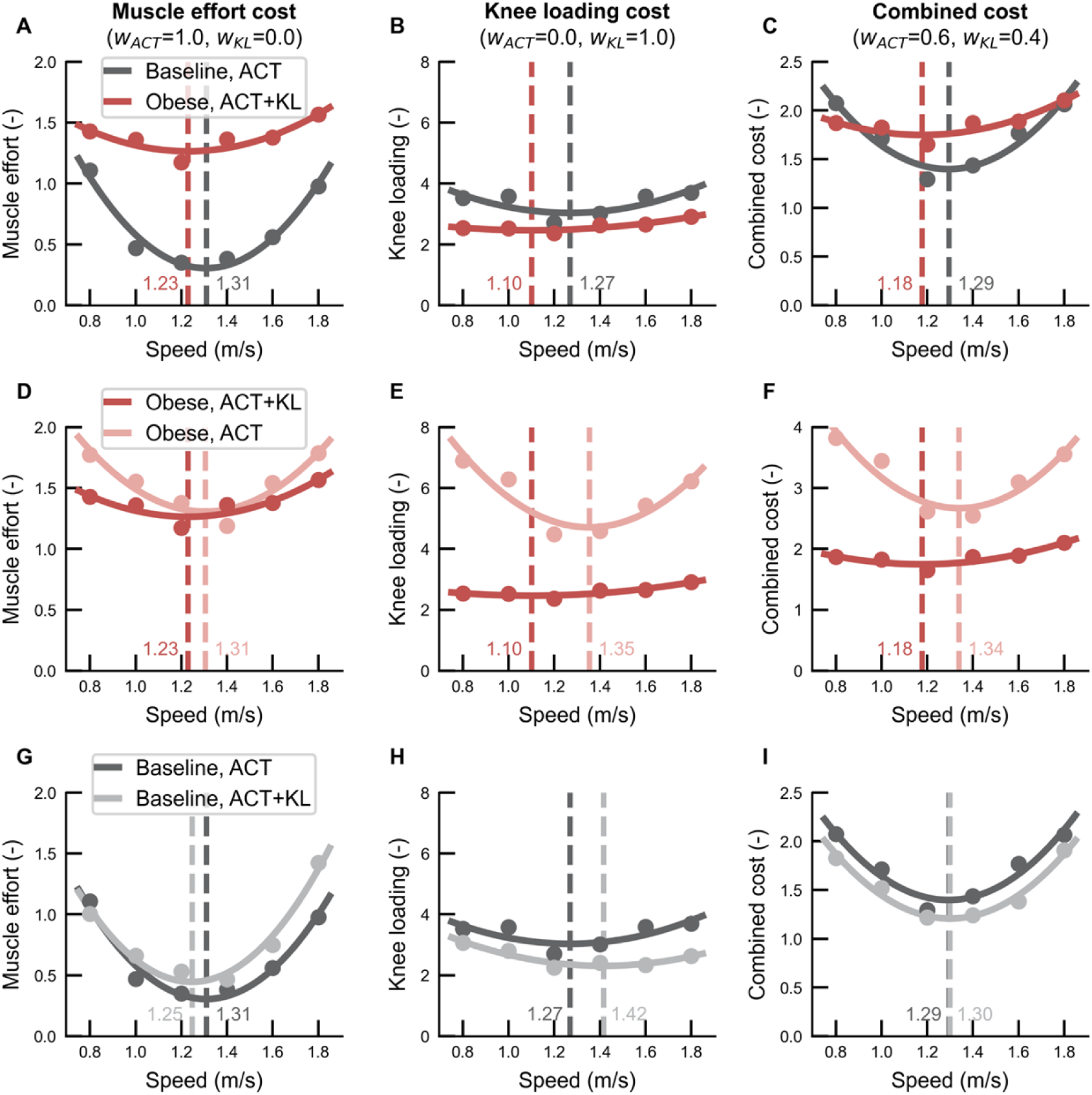
Optimal walking speeds across cost evaluations for the baseline and nominal obese models. **A–C**. Baseline model optimized for muscle effort only (ACT; *w*_ACT_ = 1.0, *w*_KL_ = 0.0) and nominal obese model optimized with the selected combined cost (ACT+KL; *w*_ACT_ = 0.6, *w*_KL_ = 0.4). Optimal speeds minimizing (A) muscle effort cost, (B) knee loading cost, and (C) combined cost. **D–F**. Obese model optimized for muscle effort only (ACT) and for the combined cost (ACT+KL). **G–I**. Baseline model optimized for muscle effort only (ACT) and for the combined cost (ACT+KL). Vertical dashed lines indicate optimal speeds.

The obese model produced shorter step lengths than the baseline model across walking speeds when optimized with the selected combined objective (Fig 4), consistent with experimental observations in obesity [11,28]. In contrast, when the obese model was optimized for muscle effort only (Obese, ACT), step lengths were similar to or larger than those of the baseline model (Fig 4A, 4B). When the baseline model was optimized using the combined objective (Baseline, ACT+KL), step lengths were also reduced relative to the muscle-effort-only condition (Fig 4C).

**Fig 4.**
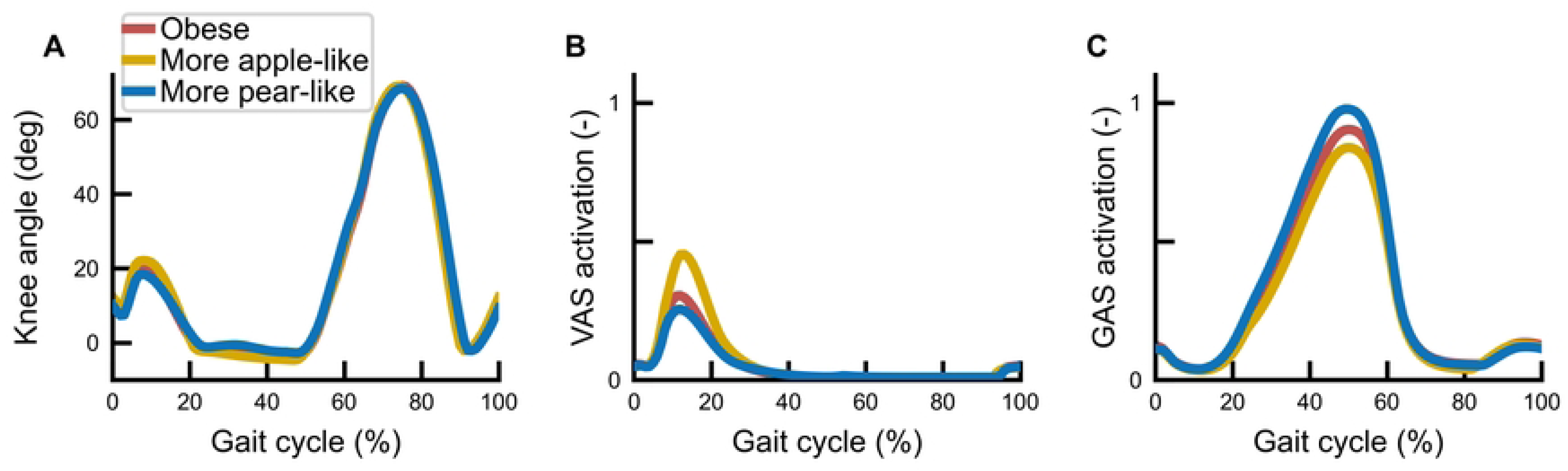
Step length across walking speeds for baseline and nominal obese models under different objective weightings. **A**. Baseline and nominal obese models optimized with their selected costs (Baseline, ACT; Obese, ACT+KL). **B**. Obese model optimized with the combined cost (Obese, ACT+KL) and for muscle effort only (Obese, ACT). **C**. Baseline model optimized for muscle effort only (Baseline, ACT) and with the combined cost (Baseline, ACT+KL).

### Body mass distribution modestly influences knee kinematics and knee muscle activation

Differences in body mass distribution produced subtle but systematic changes in gait biomechanics in the obese model variants (Fig 5). The more apple-like model exhibited slightly greater knee flexion and increased vastii activation, accompanied by slightly reduced gastrocnemius activation relative to the nominal obese model. The more pear-like model showed the opposite trends, with slightly reduced knee flexion and vastii activation and increased gastrocnemius activation.

**Fig 5.**
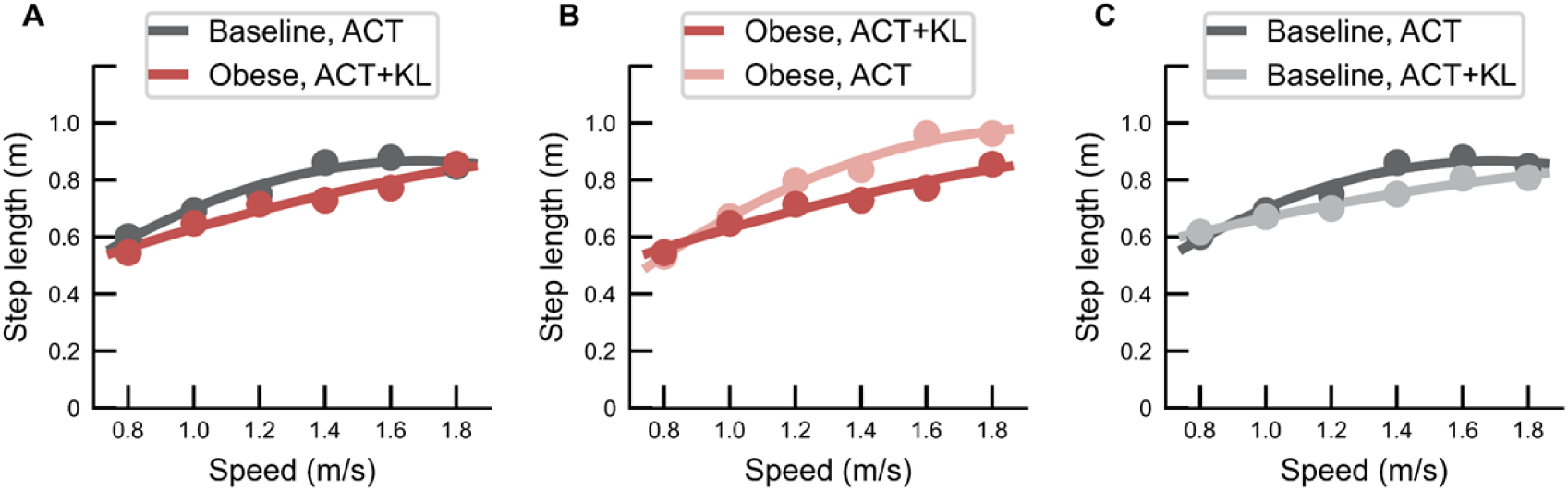
Gait biomechanics at 1.5 m/s for obese models with different body mass distributions. Nominal obese, more apple-like, and more pear-like models were simulated using the selected combined cost (ACT+KL; *w*_ACT_ = 0.6, *w*_KL_ = 0.4). **A**. Knee angle. **B**. Vastii activation. **C**. Gastrocnemius activation.

## Discussion

This study used predictive neuromechanical simulation to examine the mechanisms underlying gait adaptations associated with obesity. Obesity-related physiological changes alone did not reproduce experimentally observed gait features, but incorporating an objective that penalizes both muscle effort and knee loading generated key characteristics of obese gait, including reduced early-stance knee flexion (“stiff-knee” gait), altered muscle coordination, slower preferred walking speeds, and shorter step lengths. These results suggest that obesity-related gait patterns emerge from the interaction between altered physiology and movement objectives that balance energetic and joint loading demands.

### Mechanistic interpretation of main findings on obesity-related gait adaptations

Obesity-related gait adaptations in our simulations emerged from the interaction between altered musculoskeletal physiology and movement objectives that regulate knee loading. The non-obese baseline model reproduced typical healthy gait when optimized solely to minimize muscle effort, consistent with prior predictive simulation studies of human walking [27]. However, incorporating obesity-related changes in body mass distribution and muscle strength alone did not reproduce the reduced early-stance knee flexion observed in obesity [14,29]. Conversely, modifying the objective alone did not produce obese-like gait behaviors in the baseline model (e.g., optimal walking speeds of Baseline, ACT+KL differed from those of Obese, ACT+KL). Only when obesity-related physiological changes were paired with an objective that penalizes both muscle effort and knee loading did the model reproduce multiple experimentally observed features, including stiff-knee gait, altered muscle coordination, slower optimal walking speeds, and shorter step lengths [14,28,30]. Together, these findings suggest that characteristic gait differences associated with obesity arise from the interaction between altered musculoskeletal mechanics and coordination strategies that reduce knee extensor demand under increased loading conditions.

The reduced early-stance knee flexion in the obese model optimized with the combined objective was accompanied by decreased vastii (VAS) activation and increased reliance on ankle plantarflexors (GAS and SOL), consistent with experimental observations of distal redistribution of muscle demand in individuals with obesity [5,12,16,31]. Reduced knee flexion decreases the knee extensor moment required during weight acceptance, thereby lowering quadriceps demand [14,29]. The altered coordination pattern also influenced the distribution of tibiofemoral contact force contributions (Fig 6), where reduced VAS contribution was associated with lower accumulated knee load per distance traveled when normalized to body weight compared with the non-obese model, although absolute knee loading remained higher due to greater body mass. This result is consistent with reports that individuals with obesity exhibit lower knee contact forces relative to body mass during walking, despite greater absolute loading due to increased body weight [15,32–34]. Together, these findings suggest that the stiff-knee gait pattern and associated muscle coordination reflect a coordination strategy that reduces knee extensor demand and modulates knee joint loading under increased body mass.

**Fig 6.**
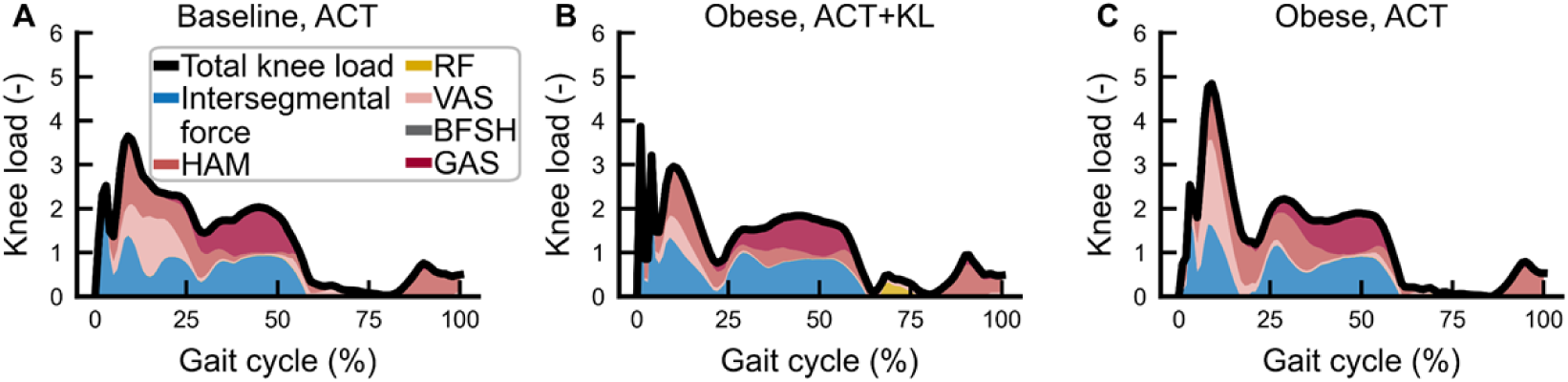
Normalized knee contact force during walking at 1.5 m/s across the gait cycle. A. Baseline, ACT (*w*_ACT_ = 1.0, *w*_KL_ = 0.0). B. Obese, ACT+KL (*w*_ACT_ = 0.6, *w*_KL_ = 0.4). C. Obese, ACT (*w*_ACT_ = 1.0, *w*_KL_ = 0.0). Knee contact force was normalized by body weight (BW). Stacked areas represent contributions from intersegmental dynamics and selected muscles (RF, VAS, HAM, GAS, BFSH). The black line indicates total knee contact force.

The combined objective also influenced preferred walking speed and step length. The obese model exhibited slower optimal walking speeds than the baseline model when evaluated using muscle effort, knee loading, and combined costs, consistent with experimentally observed reductions in preferred walking speed in individuals with obesity [30]. Shorter step lengths also emerged across walking speeds without explicit constraints on step length, suggesting that these spatiotemporal adaptations arise naturally from the same coordination strategy. This result is consistent with previous studies showing that shorter step lengths can reduce cumulative knee joint loading per distance traveled, whereas longer steps may increase energetic cost [28,35]. Together, these findings suggest that slower walking speeds and shorter step lengths may reflect coordinated adaptations that reduce cumulative knee loading under increased body mass.

Differences in body mass distribution produced modest but systematic variations in gait biomechanics. Compared with the nominal obese model, the more apple-like model showed slightly greater knee flexion and vastii activation, whereas the more pear-like model exhibited slightly greater gastrocnemius activation and slightly reduced knee flexion. Waist and hip circumferences for these variants were varied by *±*1 standard deviation from the nominal obese values based on population statistics, and additional exploratory simulations (not reported) using *±*2 standard deviations produced similar qualitative trends with only modest changes in gait outcomes. These results suggest that segment mass distribution influences joint moment demands and muscle coordination, but the magnitude of these effects was small relative to those associated with increased total body mass. Overall, the findings indicate that increased body mass is the primary driver of the observed gait adaptations, while body mass distribution provides a secondary modulation of coordination patterns.

### Clinical implications for knee joint loading and osteoarthritis risk

Obesity is a well-established risk factor for knee osteoarthritis (OA), due in part to increased mechanical loading associated with greater body mass [33,34]. Our simulations suggest that obesity-related gait adaptations may influence not only the magnitude but also the distribution of knee joint loading through altered coordination patterns. The reduced knee flexion and decreased vastii activation observed in the obese model were associated with lower accumulated knee load per distance traveled when normalized to body weight, even though absolute knee loading increased with greater body mass, consistent with reports that individuals with obesity exhibit lower knee contact forces relative to body mass during walking [14,15,32]. However, lower normalized loading does not necessarily imply reduced OA risk, because reduced knee flexion excursion during weight acceptance may alter tibiofemoral contact mechanics and shift loading toward more localized joint regions [36,37]. These findings suggest that obesity-related gait adaptations may regulate knee loading under increased body mass while still influencing mechanical factors associated with OA development and progression.

Our results also suggest that reducing body mass alone may not fully restore normal-weight-like gait coordination, because the obese and baseline models were associated with different movement objectives that regulate knee loading. Applying the combined objective to the baseline model produced differences in stance phase knee flexion (Fig 1A), optimal walking speed (Fig 3G–I), and step length (Fig 4C), indicating that coordination strategies that prioritize reduced knee loading can alter multiple gait characteristics even without increased body mass. Previous studies have reported improvements in gait mechanics following weight loss, particularly after bariatric surgery, but these studies typically compare pre- and post-weight-loss conditions within individuals rather than against normal-weight controls [38–40]. Interventions for obesity-related gait impairment may benefit from addressing not only body mass reduction but also coordination patterns influencing knee loading.

### Predictive neuromechanical simulation in the study of abnormal gait

Predictive neuromechanical simulation was particularly useful in this study because it allowed us to identify which modeling components were necessary to reproduce obesity-related gait. When obesity-related musculoskeletal changes (i.e., increased body mass and altered muscle strength) were considered in isolation, the simulations did not reproduce characteristic features of obese gait such as reduced early-stance knee flexion and shorter step length. We therefore considered alternative movement objectives that could plausibly influence coordination under increased body mass, and knee loading was a natural candidate because of its direct mechanical relevance to knee burden and OA risk in obesity. By systematically varying the weighting between muscle effort and knee loading (Fig 1), we identified an objective setting that reproduced the obese gait features of interest. Once this setting was established, the framework enabled deeper analysis of detailed gait biomechanics and walking speed dependence, as well as follow-up “what-if” investigations such as the effect of body mass distribution. More broadly, predictive simulation requires identifying physiological parameters, control structure, and objective functions that together reproduce observed gait patterns. In this study, inclusion of the knee-loading cost term was critical for reproducing obesity-related gait adaptations.

Objective functions in predictive walking simulations have most commonly emphasized muscle effort or metabolic energy expenditure, often together with task-related constraints needed to produce stable gait [21,22,27]. The knee-loading term in this study was motivated by the mechanical relevance of tibiofemoral loading in obesity but is less commonly used in predictive walking simulations. Including this term consistently altered the predicted gait pattern, most notably by producing shorter step lengths than muscle-effort-only optimization in both the obese model and the cross-objective baseline simulations (Fig 4B, 4C), consistent with prior simulation work that minimized knee joint contact force [41]. Because shorter step length is common in many abnormal gait patterns, these results suggest that knee-loading objectives may be worth investigating more broadly in predictive simulations when regulation of joint loading is a plausible component of the adaptation.

### Limitations and future directions

Several limitations should be considered when interpreting these findings. First, the present model remains a simplified representation of obesity-related gait: the 2-D sagittal-plane formulation cannot capture frontal-plane mechanics or balance-related adaptations, and the omission of trunk muscles and a toe joint likely influenced some predicted hip and ankle mechanics. In addition, obesity-related physiology was represented primarily through changes in body mass distribution and muscle strength, whereas other potentially relevant factors, including altered sensorimotor function, sex-specific morphology, and age-related muscle deficits, were not incorporated. Some of these omissions were intentional design choices to isolate the effects of increased body mass and muscle strength from neurological or age-related changes that vary substantially across individuals. Second, the objective formulation was limited to muscle effort and knee loading, so the identified solution should be interpreted as a plausible explanation for obesity-related gait rather than a unique one. Third, predictive simulations can admit multiple local minima, meaning that the identified solution may not represent the global optimum and alternative coordination patterns could produce comparable performance. To mitigate this risk, we performed multiple optimizations for each condition using different random seeds and optimization settings, including chained runs, and selected the best-performing solutions following our prior practice [27,42]. Finally, this study considered only a male musculoskeletal model, because extending the framework to females would require a separate baseline model with sex-specific anthropometric and musculoskeletal parameters that is not currently available in our framework. Future work could extend this approach to female models, as well as to more complete 3-D and subject-specific representations, a broader range of locomotor tasks such as slopes, stairs, turns, and perturbations, and additional candidate physiological and control mechanisms that may contribute to abnormal gait.

In conclusion, predictive neuromechanical simulation identified a mechanistic explanation for characteristic gait adaptations associated with obesity and their potential relevance to knee osteoarthritis. The findings indicate that obesity-related gait patterns emerge from the interaction between altered musculoskeletal physiology and coordination strategies that balance muscle effort and knee joint loading. Incorporating a knee-loading objective was necessary to reproduce reduced early-stance knee flexion, altered muscle coordination, slower optimal walking speeds, and shorter step lengths observed experimentally, suggesting that these adaptations may reflect strategies to regulate tibiofemoral loading under increased body mass. Although such adaptations may reduce knee load relative to body weight, absolute knee loading still increases and coordination changes may influence the distribution and cumulative exposure of mechanical factors implicated in OA development and progression. Variations in body mass distribution produced consistent but moderate effects on gait, indicating that total body mass is the primary driver of these changes. Together, these results demonstrate the utility of predictive neuromechanical simulation for identifying candidate mechanisms linking obesity, gait biomechanics, and knee joint loading, and provide a framework for investigating how physiological and coordination changes may influence joint health.

## Materials and Methods

We used a predictive neuromechanical simulation framework previously developed and validated in our prior studies [25,42] to investigate gait adaptations associated with obesity. A baseline model representing a healthy adult (1.8 m, 80 kg) was modified to incorporate obesity-related changes in segmental mass, inertia, and muscle strength. Control parameters were optimized to generate stable walking that minimized muscle effort and knee loading. Simulations were conducted across walking speeds and body mass distributions to evaluate resulting gait biomechanics. All simulations were implemented in MATLAB (R2023a) using Simulink and Simscape Multibody to model the forward dynamic interaction between the musculoskeletal system and reflex-based controller.

### Neuromechanical simulation framework and baseline model

The predictive neuromechanical simulation framework consists of a closed-loop interaction between a musculoskeletal system, a reflex-based locomotion controller, and an optimization process that adjusts control parameters to generate stable walking (Fig 7). Within the forward dynamic neuromechanical simulation, rigid-body skeletal dynamics and musculotendon actuators interact with the neural controller through sensory feedback pathways, such that movement emerges from the coupled dynamics of muscle forces, segmental mechanics, and foot–ground contact. Muscle stimulation signals generated by the reflex-based controller activate muscles, while biologically plausible sensory signals describing body state are fed back to the controller to regulate locomotion. The resulting gait patterns are evaluated using an objective function, and controller parameters are iteratively updated through parameter optimization to produce steady walking behaviors that minimize the objective function value. This forward dynamic formulation allows gait to emerge from the interaction between biomechanics and neural control, without tracking or prescribing experimental motion data (e.g., kinematic trajectories). The framework is adapted from our previous studies [25,42].

**Fig 7.**
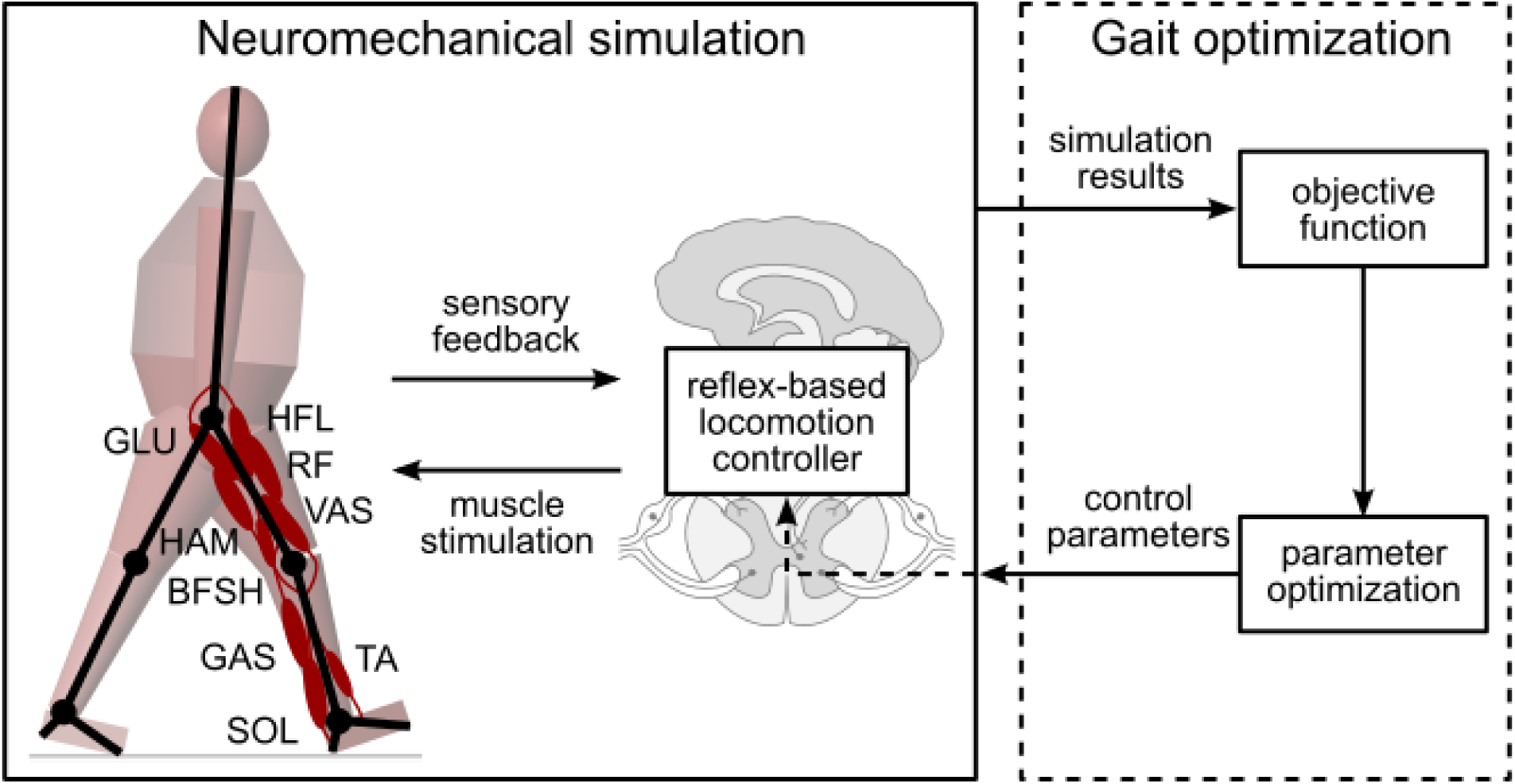
Predictive neuromechanical simulation framework. Schematic illustrating the closed-loop interaction between the musculoskeletal system, reflex-based neural controller, and optimization process. Sensory feedback signals regulate muscle stimulation to generate walking through forward dynamic simulation, while controller parameters are iteratively updated to minimize the objective function.

The baseline musculoskeletal model is a 2D sagittal-plane representation of a healthy adult male (height 1.8 m, mass 80 kg). The model consists of seven rigid-body segments: a single segment representing the head, arms, and trunk, and three segments per leg representing the thigh, shank, and foot. These segments are connected by hinge joints, resulting in six internal degrees of freedom corresponding to the hip, knee, and ankle joints of each leg. Movement is generated by musculotendon actuators modeled using Hill-type muscle models, which capture the dependence of muscle force on muscle length and contraction velocity observed in biological muscle [43]. Each leg is actuated by nine musculotendon units representing major lower-limb muscle groups: hip flexors (HFL), glutei (GLU), hamstrings (HAM), rectus femoris (RF), vastii (VAS), short head of the biceps femoris (BFSH), gastrocnemius (GAS), soleus (SOL), and tibialis anterior (TA). Muscle-specific parameters include maximum isometric force, optimal fiber length, maximum contraction velocity, and tendon slack length for each musculotendon unit. Muscle forces are converted into joint torques through variable moment arms that depend on joint configuration. These parameter values and musculoskeletal architecture are based on physiological measurements reported in the literature and are consistent with widely used models in predictive neuromechanical simulation studies and our prior work [27,42,43]. Muscle activations are generated by a reflex-based controller that encodes key functions required for stable human locomotion using feedback pathways that primarily represent spinal reflex circuits [25]. The controller uses physiologically plausible sensory signals related to muscle force (Golgi tendon organs), muscle fiber length and velocity (muscle spindles), ground contact and leg loading (cutaneous receptors and load-sensitive afferents), and trunk orientation relative to gravity (vestibular pathways) to coordinate essential locomotor functions, including compliant stance leg support, swing leg placement for balance, and stabilization of trunk posture. The full controller includes 64 tunable parameters, such as reflex gains and length offsets. When optimized to minimize muscle activation or metabolic energy expenditure, reflex-based controllers of this class produce stable, human-like gait across a range of walking speeds and environmental conditions [25], including simulations incorporating age-related musculoskeletal changes [27]. Full details of the control architecture are provided in the original publications [25].

### Obesity-related musculoskeletal model modifications

We modified the baseline musculoskeletal model to represent individuals with obesity by adjusting segmental mass, inertia, and muscle strength. The reflex-based controller structure was kept unchanged, while control parameters were optimized separately for each model using an objective formulation. Segment properties were estimated using a data-driven anthropometric scaling model developed in our recent work [44], which was specifically designed to account for variation in body composition observed in individuals with obesity and predicts segment mass distribution from height, body mass, sex, and body shape descriptors based on large population datasets (ANSUR II [45] and NHANES [46]). The nominal obese model represented an adult male with height 1.8 m and body mass 140 kg (BMI 43 kg/m^2^). To examine the influence of body mass distribution, additional obese variants with more apple-like (greater upper-body mass concentration) and more pear-like (greater lower-body mass concentration) shapes were generated by varying waist-to-hip ratio using the same anthropometric scaling model [44]. Waist and hip circumferences for these variants were set to one standard deviation above or below the nominal obese values based on population statistics from the ANSUR II database [45]. Muscle adaptations associated with obesity were represented by increasing maximum isometric muscle force by 20% to reflect hypertrophy in weight-bearing muscles subjected to chronic overload, while still resulting in reduced strength relative to body mass [7,8]. Key anthropometric and musculoskeletal parameters for all model variants are summarized in Table 2.

**Table 2.**
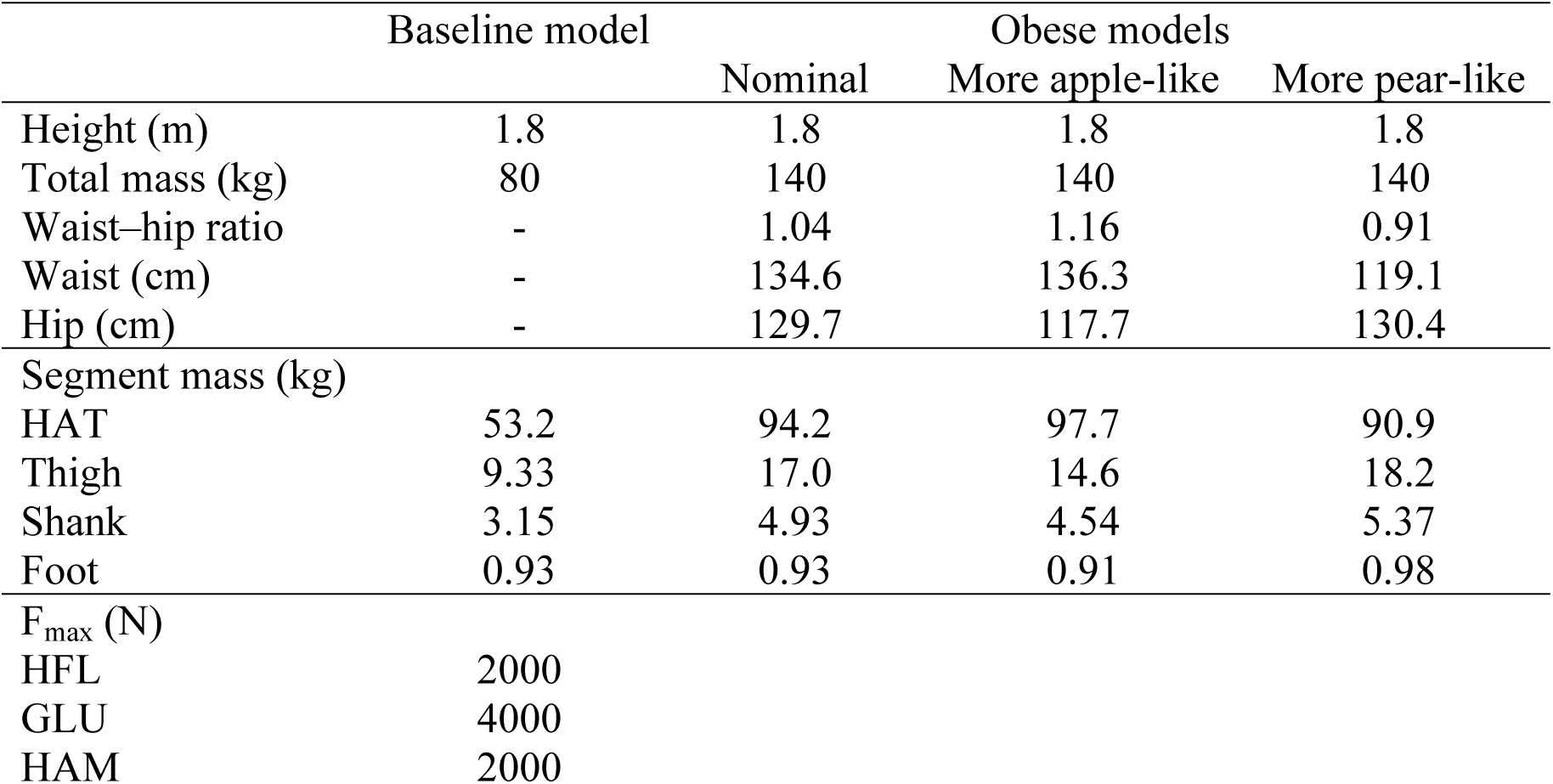
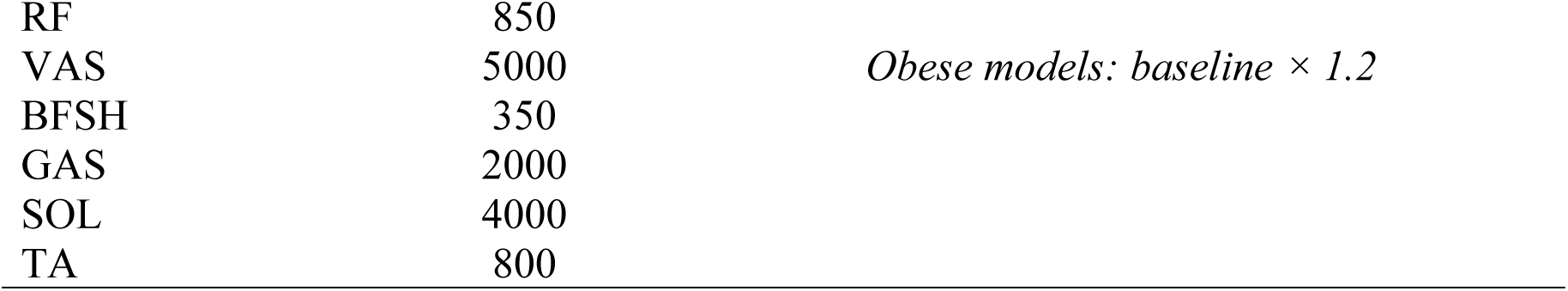
Anthropometric characteristics, segment masses, and maximum isometric muscle forces (Fmax) for the baseline and obesity model variants. Waist and hip values denote circumferences.

|  | Baseline model | Obese models |  |  |
| --- | --- | --- | --- | --- |
|  |  | Nominal | More apple-like | More pear-like |
| Height (m) | 1.8 | 1.8 | 1.8 | 1.8 |
| Total mass (kg) | 80 | 140 | 140 | 140 |
| Waist–hip ratio | - | 1.04 | 1.16 | 0.91 |
| Waist (cm) | - | 134.6 | 136.3 | 119.1 |
| Hip (cm) | - | 129.7 | 117.7 | 130.4 |
| Segment mass (kg) |  |  |  |  |
| HAT | 53.2 | 94.2 | 97.7 | 90.9 |
| Thigh | 9.33 | 17.0 | 14.6 | 18.2 |
| Shank | 3.15 | 4.93 | 4.54 | 5.37 |
| Foot | 0.93 | 0.93 | 0.91 | 0.98 |
| F<sub>max</sub> (N) |  |  |  |  |
| HFL | 2000 |  |  |  |
| GLU | 4000 |  |  |  |
| HAM | 2000 |  |  |  |

|  |  |  |
| --- | --- | --- |
| RF | 850 | *Obese models: baseline $\times 1.2$* |
| VAS | 5000 |  |
| BFSH | 350 |  |
| GAS | 2000 |  |
| SOL | 4000 |  |
| TA | 800 |  |

### Control parameter optimization for gait prediction

Control parameter optimization was performed to generate predictive walking simulations for each musculoskeletal model and prescribed walking speed (Fig 7). For each model variant, 64 parameters of the reflex-based controller, together with 7 parameters defining the initial pose, were optimized to produce stable and steady gait without tracking experimental kinematics. Optimization was performed using the covariance matrix adaptation evolution strategy (CMA-ES) [47], which iteratively improves candidate solutions based solely on the ranking of objective function values (without requiring gradient information or smoothness of the cost landscape).

The objective formulation guided the model toward generating stable walking at the prescribed speed while minimizing muscle effort and knee joint loading. We adopted a staged cost structure [25] that first penalizes simulations that fall, followed by simulations that fail to achieve steady gait or violate key biomechanical constraints such as target speed, joint limits, or trunk posture. The staged cost is defined as

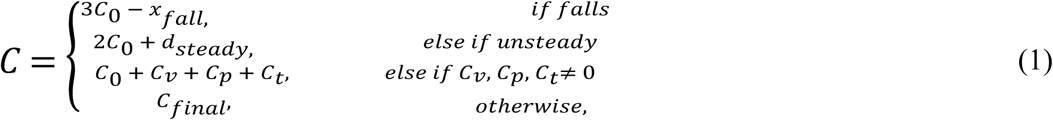

where *C_0_* = 10^4^ is a large constant used to separate the four stages of evaluation*, x_fall_* is the distance walked before falling, and *d_steady_* quantifies step-to-step kinematic variations. The terms *C_v_, C_p_* and *C_t_* penalize deviations from the prescribed walking speed, violations of joint limits through passive joint torques, and excessive trunk lean, respectively [42]. Simulations that satisfy these constraints are evaluated using the final performance objective, which combines muscle effort and knee-loading terms:

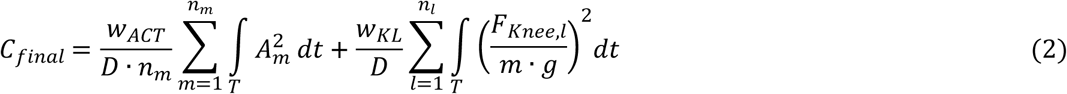

where *D* is the distance traveled, *n_m_* is the number of muscles, *A_m_* is the activation of muscle *m*, *F*_knee*,l*_ is the knee contact force in leg *l* (i.e., the compressive component of the tibiofemoral joint contact force aligned with the tibial axis, arising from intersegmental dynamics and muscle forces [48]), *m* is the body mass, *g* is gravitational acceleration, *T* is the time duration of steady walking used for cost evaluation, and *w*_ACT_ and *w*_KL_ are weighting factors for muscle effort and knee loading. The muscle effort term reflects the tendency of the neuromuscular system to favor coordination patterns that reduce overall muscle effort, a formulation widely used in musculoskeletal simulations [27], while the knee-loading term was introduced to evaluate whether gait adaptations associated with obesity may reflect strategies that reduce excessive tibiofemoral joint loading [14,15]. Both terms were normalized by walking distance to reflect performance over a given travel distance.

Each predictive walking simulation was obtained through an independent CMA-ES optimization [47] in the 71-dimensional control parameter space. To improve convergence robustness, optimizations were initialized using solutions from nearby walking speeds or related model variants. For obese models, segment masses and corresponding muscle strengths were gradually increased from baseline values to guide the search toward stable walking solutions. CMA-ES was run with a population size of 32 candidate solutions per generation for 800 generations. A typical optimization required approximately 8 hours on the Boston University Shared Computing Cluster (SCC) using 32 CPU cores in parallel.

### Simulation experiments and gait analysis

We performed simulation experiments to evaluate predicted gait biomechanics under different objective weightings, walking speeds, and body mass distributions. We first identified the weighting between muscle effort and knee loading that best reproduces experimentally observed knee kinematics, then generated predictive simulations across walking speeds for baseline and obese models, and finally evaluated the influence of body mass distribution using more apple-like and more pear-like model variants.

#### Identification of objective weightings for muscle effort and knee loading

We simulated walking for the nominal obese model across a range of objective weightings for muscle effort and knee loading to identify values that best reproduce experimentally observed knee kinematics. There were a total of 11 weight combinations, where weights *w*_ACT_ and *w*_KL_ were varied from 0 to 1 in increments of 0.1 while maintaining *w*_ACT_ + *w*_KL_ = 1. Simulations were performed at a target walking speed of 1.5 m/s, matching the speed reported in the reference experimental data [15]. Weightings were selected to reproduce experimentally observed knee kinematics associated with obesity, specifically reduced knee flexion during early stance (0–30% of the gait cycle) [15]. Agreement between simulated and experimental knee angle trajectories was quantified using the root mean squared error (RMSE) of knee flexion angle during early stance. We also examined the resulting joint angles, joint moments, ground reaction forces, and muscle activations. The resulting weight sets for each of the baseline and nominal obese models were used for all remaining simulations.

#### Baseline and obese gait simulations across walking speeds

Using the selected objective weighting described above, we generated predictive walking simulations for both the baseline and nominal obese models across a range of walking speeds. Each model was optimized for speeds (0.8, 1.0, 1.2, 1.4, 1.6, 1.8 m/s). We analyzed muscle effort, knee loading, and step length across walking speeds.

#### Body shape variation: more apple-like and more pear-like models

We simulated walking for the more apple-like and more pear-like obese models to evaluate the influence of body mass distribution on gait biomechanics. We examined differences in predicted gait at a walking speed of 1.5 m/s.

## Acknowledgements

The authors acknowledge the Boston University Shared Computing Cluster (SCC), which provided 59,600 hours of computing resources for this project.

## Data availability

All custom code used to process the data from simulations and generate the figures in this study (including Python scripts) is openly available on https://github.com/cwchoi4105/Predictive-Neuromechanical-Simulation-Obesity or https://zenodo.org/records/20492556. The repository also contains simulation results such as time series joint angle, moment, muscle activation and knee force derived from the simulations, and these files allow users to reproduce the main figures.

## CRediT authorship contribution statement

Chi-Whan Choi, Simone V. Gill, and Seungmoon Song: Conceptualization. Chi-Whan Choi, Vincent Ton and Seungmoon Song: Methodology, Software, Visualization. Chi-Whan Choi: Writing – Original draft preparation. Chi-Whan Choi, Vincent Ton, Simone V. Gill and Seungmoon Song: Writing – Reviewing and editing. All authors approved the final manuscript before submission.

## Supporting information

### Supplementary Video

Walking simulation at 1.5 m/s in Baseline, ACT (*w*_ACT_ = 1.0, *w*_KL_ = 0.0), Obese, ACT (*w*_ACT_ = 1.0, *w*_KL_ = 0.0) and Obese, ACT+KL (*w*_ACT_ = 0.6, *w*_KL_ = 0.4) models across the gait cycle.

